# An intein-mediated split–nCas9 system for base editing in plants

**DOI:** 10.1101/2021.10.08.463716

**Authors:** Guoliang Yuan, Haiwei Lu, Md Mahmudul Hassan, Yang Liu, Yi Li, Paul E. Abraham, Gerald A. Tuskan, Xiaohan Yang

**Author notes:** Corresponding authors: Gerald A. Tuskan; Xiaohan Yang. Department of Academic Education, Central Community College – Hastings, Hastings, NE 68902, USA. **Disclosure:** This manuscript has been authored by UT-Battelle, LLC under Contract No. DE-AC05-00OR22725 with the U.S. Department of Energy. The United States Government retains and the publisher, by accepting the article for publication, acknowledges that the United States Government retains a non-exclusive, paid-up, irrevocable, worldwide license to publish or reproduce the published form of this manuscript, or allow others to do so, for United States Government purposes. The Department of Energy will provide public access to these results of federally sponsored research in accordance with the DOE Public Access Plan (http://energy.gov/downloads/doe-public-access-plan).

## Abstract

Virus-assisted delivery of the clustered regularly interspaced short palindromic (CRISPR)/CRISPR-associated (Cas) system represents a promising approach for editing plant genomes. However, the relatively large size of the CRISPR/Cas9 system is challenging to package into viral vectors with confined packaging capacity. To address this technical challenge, we developed a strategy that splits the required CRISPR-Cas9 components across a dual-vector system in which CRISPR-Cas reassembles into an active form following co-infection to achieve targeted genome editing in plant cells. An intein-mediated split system was adapted and optimized in plant cells by successfully demonstrating split-eYGFPuv expression. Using a plant-based biosensor, we demonstrated for the first time that the split-SpnCas9 is capable of inducing efficient base editing in plant cells and identified several valid split sites for future biodesign strategies. Overall, this strategy provides new opportunities to bridge different CRISPR/Cas9 tools including base editor, prime editor, and CRISPR activation with virus-mediated gene editing.

## Introduction

The programmable clustered regulatory interspaced short palindromic repeat (CRISPR) and CRISPR-associated protein (CRISPR/Cas) technology has revolutionized plant genome editing. However, there are still important limitations to consider that currently impede the commercial applications of CRISPR in agriculture, particularly the presence of transgenes. ^1^ In plants, virus- and nanoparticle-mediated deliveries of a CRISPR/Cas system are two promising methods to create transgene-free targeted mutants without requiring the time consuming tissue culture process. ^2, 3^ Virus-assisted delivery of CRISRP/Cas systems has been successfully demonstrated in plants as a method to circumvent conventional tissue culture-based methods; however, only until recently, these viruses could only be used for sgRNA delivery. ^4, 5^ Strategies to overcome limited cargo capacities in viruses are crucial for next-generation virus-assisted genome engineering. Most recently, using a negative-stranded RNA virus with a cargo capacity large enough for an entire CRISPR-Cas9 cassette, targeted DNA-free genome editing has been achieved in tobacco. ^6^ Unfortunately, because each virus species has inherent host range constraints, this viral delivery method will have limited applicability across plant species. As such, there is a crucial need for strategies that can accommodate different viral-delivery systems.

A promising approach to overcome technical challenges associated with virus mediated CRISPR/Cas genome editing is using a split-protein. An intein-mediated split-*Sreptococcus pyogenes* Cas9 (SpCas9) system was recently demonstrated in human cells, whereby the SpCas9 nuclease protein coding system was functionally split into two inactivate fragments across a dual-vector system, delivered, and its activity reconstituted efficiently in cells via co-expression. ^7^ In plant cells, a split *Staphylococcus aureus* Cas9 (SaCas9) is able to induce targeted mutagenesis for transgene-free genome editing. ^8^ To date, the reported split sites of Cas9 are still limited, and much less is known about how to effectively split the most commonly used SpCas9 protein used for gene editing in plants. Therefore, we developed an intein-mediated split SpCas9 nickase (SpnCas9, D10A) for base editing in plant protoplasts that functions with high efficiency and comparable performance to wild-type full-length SpnCas9.

## Results and Discussion

We used one well-characterized split intein, derived from *Npu*DnaE ^9^ for splitting SpnCas9. Protein splicing elements (called “inteins”) allow the coding sequence of a target protein to be split into two inactive fragments and reconstituted post-translationally (Figure 1a). To improve gene expression and increase translational efficiency, *Arabidopsis* codon optimization of the *Npu*DnaE intein was performed using the online codon optimization tool, ExpOptimizer, provided by NovoPro Bioscience (Shanghai, China). In general, the coding sequence of a target gene was split into an N-terminal fragment (Gene^N^) and a C-terminal fragment (Gene^C^), which were then cloned upstream of an N-terminal fragment of the *Npu*DnaE intein (Int^N^) and downstream of a C-terminal fragment of the *Npu*DnaE intein (Int^C^), respectively, into two different vectors (Figure 1b).

**Figure 1.**
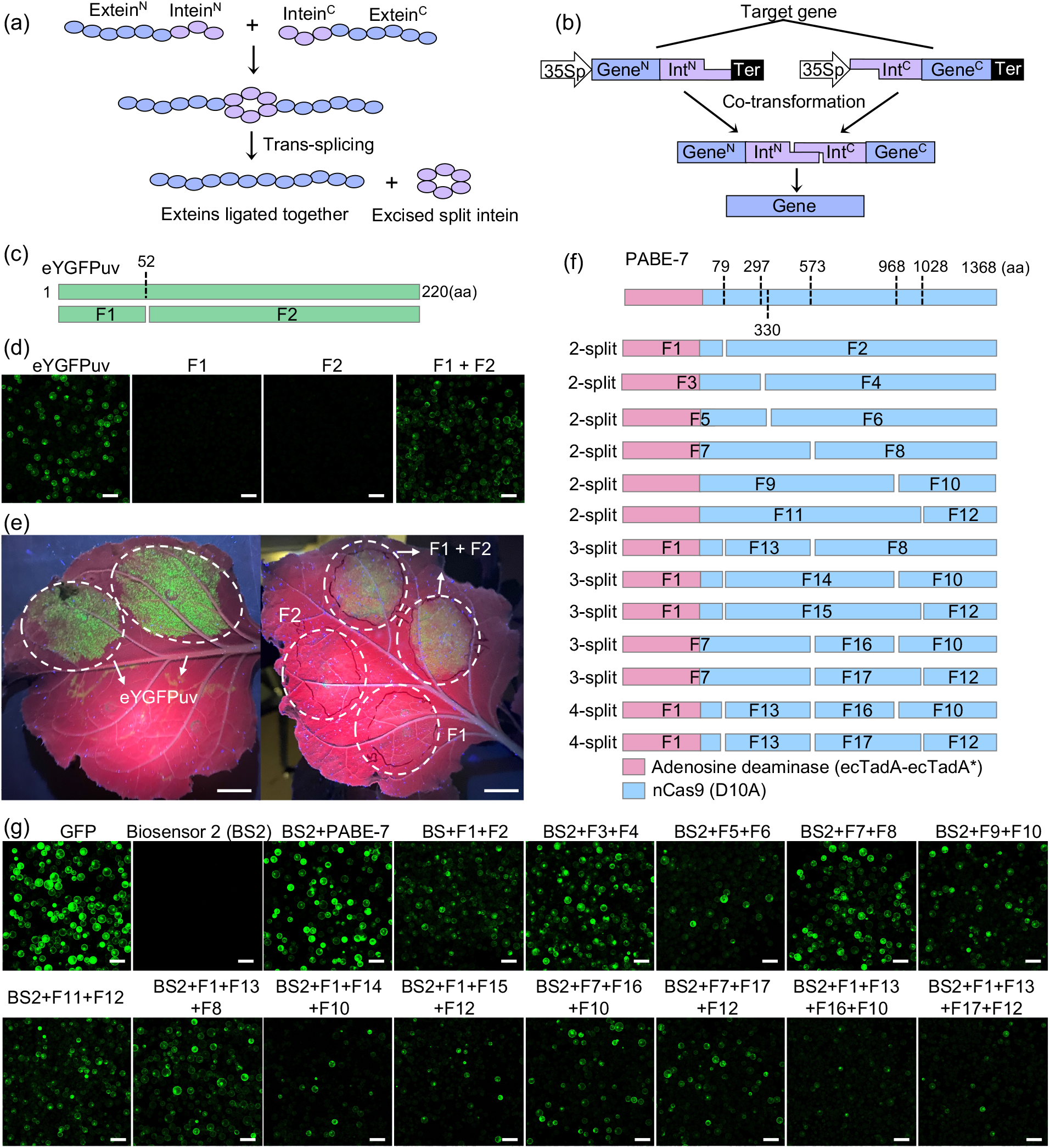
The *Npu*DnaE intein-mediated split–SpnCas9 for base editing in plant system. (a) Trans-splicing mechanism reaction by split inteins. (b) Illustration of vector design and reconstitution of a target gene. Ter, terminator. (c) Identification of potential split site in eYGFPuv (d) Transient expression of split-eYGFPuv in *Arabidopsis* protoplasts. Scale bar, 100 μm. (e) Transient expression of split-eYGFPuv in *N. benthamiana*. Scale bar, 1cm. (f) Identification of potential split site in SpnCas9. (g) Transient expression of split-SpnCas9 in *Arabidopsis* protoplasts. Scale bar, 100 μm.

To test the efficacy of the split system, an eYGFPuv reporter ^10^ was used as the target gene. The potential split site T52:C53 was identified and used to split the eYGFPuv into two fragments, F1 and F2, because it’s reported that the obligatory cysteine residue on the C-extein junction and a residue on the N-extein junction promote substantial trans-splicing activities (Figure 1c). ^9^ Then two plasmids containing F1and F2, respectively, were co-transformed into *Arabidopsis* protoplasts through PEG-mediated transformation with a 35Sp-eYGFPuv construct as a positive control. Two days post transformation, bright green fluorescence was observed under a confocal microscope in both the positive control and the protoplasts co-transformed with F1 and F2 plasmids, though the fluorescence in latter was relatively weaker, whereas no green fluorescence was detected in the protoplasts containing F1 or F2 plasmids alone (negative controls) (Figure 1d). Also, we tested split-eYGFPuv using *Agrobacterium*-mediated leaf infiltration in *Nicotiana benthamiana*. Similarly, clear green fluorescence was observed under UV light in the positive control as well as the leaf area co-infiltrated with F1 and F2 plasmids but not in the negative control (Figure 1e). These results indicate that *Npu*DnaE intein-based split system works efficiently in plant systems.

To split SpnCas9, we identified one native split site (I79:C80), one reported spilt site (E573:C574) ^7^ and created four artificial split sites (S297^C, Q330^C, K968^C and E1028^C) by inserting a cysteine on the C-extein junction. Thus, an SpnCas9 can be divided into two even fragments as illustrated in Figure 1f. Previously, to detect base editing activities in plant systems, we have developed biosensor 2 (BS2) which is composed of a *GFP* mutant harboring a premature termination codon (PTC) and a single guide RNA (sgRNA) targeting the PTC. The *GFP* mutant can be rescued by an adenine base editor (ABE) under the guidance of the sgRNA, leading to the generation of green fluorescence. ^11^ Here, the efficacy of the split SpnCas9 system was examined in *Arabidopsis* protoplasts by the co-transfection with BS2. Two days post transformation, bright green fluorescence was observed in both the positive control (35Sp:GFP) and the protoplasts co-transfected with a plant adenine base editor PABE-7 ^12^ and BS2 but not those transformed with BS2 alone, indicating that BS2 detected the base editing activity successfully in protoplasts (Figure 1g). In contrast, strong green fluorescence was also detected in the protoplasts co-expressing BS2 and intein-mediated 2-split SpnCas9 fragments, indicating that the intein-mediated split SpnCas9 successfully induced active base editing in plant cells (Figure 1g). Moreover, using the same method we examined 3-split and 4-split SpnCas9 systems containing three and four fragments, respectively, as shown in Figure 1f. Interestingly, clear GFP signal was detected in the protoplasts containing BS2 and five 3-split combinations or two 4-split combinations, while the GFP intensity was lower in comparison with those co-expressing 2-split fragments (Figure 1g). These results indicate that the intein-reconstituted split SpnCas9 system with up to four split fragments is functional in plant base editing. Furthermore, ~54% and ~35% of the cells exhibited GFP signals in the positive control and the samples with BS2-PABE-7 co-transformation, respectively (Figure 2a). In the protoplasts co-expressing BS2 and different 2-split components, approximately 9~32% of the cells exhibited GFP fluorescence (Figure 1h). GFP fluorescence was also detected in about 8%~30% and 5% of the cells in the three-fragment and four-fragment co-transformations, respectively (Figure 2a). This result indicates that the targeting efficiency of BS2 with intein-mediated split–SpnCas9 system is comparable to wild-type SpnCas9, especially in four 2-split sites (I79:C80, S297^C, E573:C574 and E1028^C) and one 3-split sites (I79:C80 and E573:C574) (Figure 2b).

**Figure 2.**
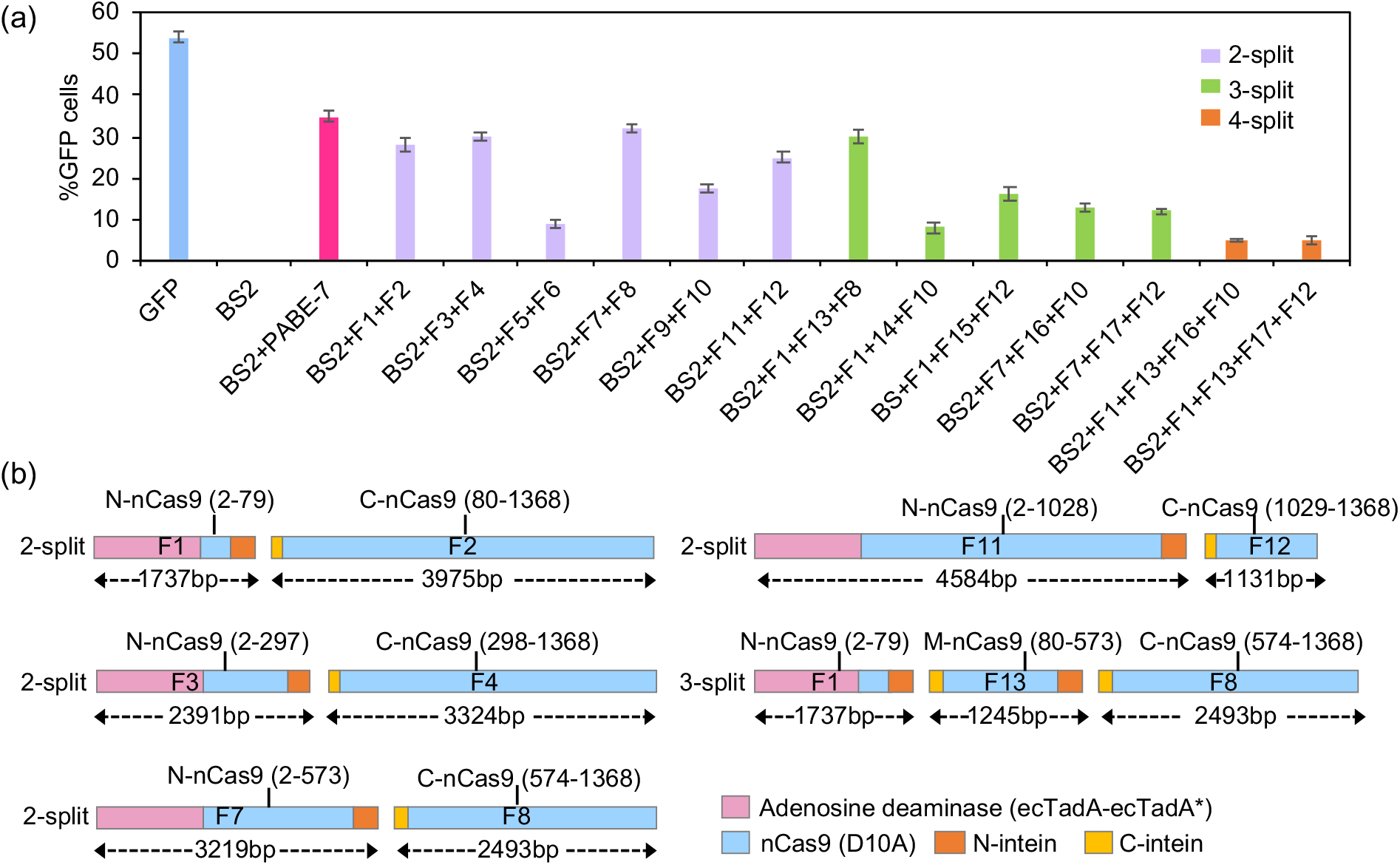
The identification of valid split sites for split–SpnCas9. (a) Statistical analysis of GFP-positive cells with different split-SpnCas9 components. All data are presented as the mean ± SE (n = 5 independent scopes). (b) Different SpnCas9 fragments with high editing efficiency. M-nCas9 (80-573), the middle fragment in 3-split.

In summary, we showed that the *Npu*DnaE intein-mediated split system functions effectively in plant systems. We also demonstrated that the base editing activity of our split-intein system is comparable to wild-type full-length SpnCas9. In addition to base editing, it has been reported that a split prime editor system could mediate endogenous base transversion and insertion in human cells. ^13^ Thus, the identification of multiple valid split sites provides more choice for split Cas9 and can be potentially used for delivery of CRISPR/Cas9 tools such as base editor, prime editor, and CRISPR interference and activation through *in-planta* transformation mediated by viruses or nanoparticles.

## Materials and methods

### Cloning

To split eYGFPuv, a gblocks containing 5’-eYGFPuv and N-terminal of NpuDnaE were synthesized from Integrated DNA Technologies IDT. The F1 fragment of eYGFPuv was assembled by cloning the gblocks and a PCR-amplified relevant fragment into an eYGFPuv vector ^10^ through NEBuilder HiFi DNA Assembly (New England BioLabs, Catalog #E5520S). Similarly, the F2 fragment of eYGFPuv was created using a gblocks containing 3’-eYGFPuv and C-terminal of NpuDnaE. To split SpnCas9, a PCR-amplified 5’-PABE-7 fragment and a gblcoks containing N-terminal of NpuDnaE were assembled into the eYGFPuv vector, generating F1, F3, F5, F7, F9 and F11. Then a PCR-amplified 3’-PABE-7 fragment and a gblcoks containing C-terminal of NpuDnaE were used to assemble F2, F4, F6, F8, F10 and F12. The construction of F13 to F17 were completed by cloning a gblocks containing N-terminal of NpuDnaE, a middle fragment of PABE-7 and a PCR-amplified C-terminal of NpuDnaE into the eYGFPuv vector. DNA sequences encoding inteins were codon optimized for *Arabidopsis* using the online codon optimization tool (ExpOptimizer) provided by NovoPro Bioscience. Positive plasmids were selected through colony PCR and verified by Sanger sequencing.

### Protoplast transformation

The isolation and transient transformation of *Arabidopsis* leaf mesophyll protoplasts were performed as described previously. ^14^

### Tobacco leaf infiltration

*Agrobacterium* strain GV3101 harboring the plasmid of interest was injected into *N. benthamiana* leaves using a syringe without a needle as described by Li. ^15^

## Author contributions

G.Y. and X.Y conceived the research. G.Y. conducted the experiments and wrote the paper. All authors revised the manuscript.

## Acknowledgements

The manuscript is supported by the Center for Bioenergy Innovation (CBI), a U.S. Department of Energy (DOE) Research Center and the Secure Ecosystem Engineering and Design (SEED) project funded by the Genomic Science Program of the U.S. Department of Energy, Office of Science, Office of Biological and Environmental Research (BER) as part of the Secure Biosystems Design Scientific Focus Area (SFA). Oak Ridge National Laboratory is managed by UT-Battelle, LLC for the U.S. Department of Energy under Contract Number DE-AC05-00OR22725.

